# Superficial Slow Rhythms Integrate Cortical Processing in Humans

**DOI:** 10.1101/202549

**Authors:** Mila Halgren, Daniel Fabo, István Ulbert, Joseph R. Madsen, Lorand Eröss, Werner K. Doyle, Orrin Devinsky, Donald Schomer, Sydney S. Cash, Eric Halgren

**Author notes:** Corresponding author: Mila Halgren - Department of Neurology, Epilepsy Division, Massachusetts General Hospital, Harvard Medical School, Boston, MA 02114, USA. Both authors contributed equally to this work (Sydney S. Cash, Eric Halgren).

## Abstract

The neocortex is composed of six anatomically and physiologically specialized layers. It has been proposed that integration of activity across cortical areas is mediated anatomically by associative connections terminating in superficial layers, and physiologically by slow cortical rhythms. However, the means through which neocortical anatomy and physiology interact to coordinate neural activity remains obscure. Using laminar microelectrode arrays in 19 human participants, we found that most EEG activity is below 10-Hz (delta/theta) and generated by superficial cortical layers during both wakefulness and sleep. Cortical surface grid, grid-laminar, and dual-laminar recordings demonstrate that these slow rhythms are synchronous within upper layers across broad cortical areas. The phase of this superficial slow activity is reset by infrequent stimuli and coupled to the amplitude of faster oscillations and neuronal firing across all layers. These findings support a primary role of superficial slow rhythms in generating the EEG and integrating cortical activity.

## Introduction

The human brain must coordinate and organize the activity of billions of neurons. Cortical oscillations, by rhythmically modulating neural activity, are a likely mechanism for accomplishing this^1,2^. Although the relationship of cortical rhythms to behavioral states has been studied for nearly a century, scientists still search for unifying principles governing neocortical oscillations. One commonly accepted principle is that slow rhythms coordinate activity across widespread neuronal pools, whereas fast rhythms mediate local processing^3,4^. Furthermore, slow and fast rhythms have been hypothesized to enact feedback and feedforward processes, respectively^5,6^. Parallel with this functional hypothesis, superficial cortical layers are anatomically structured to mediate global associative processing due to their lateral connectivity, feedback connections, and diffuse thalamocortical matrix afferents^7–10^. This allows superficial cortical layers to sample activity from many cortical areas simultaneously and modulate local processing accordingly. Synthesizing these two theories suggests that low frequency rhythms might be generated in superficial cortical layers, coordinate activity over broad cortical areas and modulate higher frequency local activity in deeper cortical laminae. However, no systematic relationship has yet been detailed between oscillatory activity and cortical layers in humans, and the relationship between cortical oscillations in specific layers and those across the surface remains unexplored.

To measure *in vivo* cortical oscillations with high spatiotemporal precision, we performed intracranial electroencephalography (iEEG) in 19 patients with medically intractable epilepsy. Recordings were for the most part spontaneous, including both sleep and waking, but also included task-related activity in 3 patients. To determine how cortical oscillations varied across cortical areas and layers, we combined recordings from macroelectrodes placed horizontally on the cortical surface (ECoG) with recordings from microelectrodes inserted vertically, or perpendicular to the cortical surface (laminar electrodes) in frontal, temporal and parietal association cortices^11^. Recording vertically and laterally simultaneously allowed us to assess the characteristics of oscillations at different depths within the grey matter, and to examine the relationships between activity in specific cortical laminae to those measured on the cortical surface in distal and proximal regions.

## Results

### Distribution of Local Field Potential gradients (LFPg) of different frequencies across layers

To determine which cortical layers generate different EEG frequencies, we measured the LFPg in multiple cortical layers simultaneously using the vertical microelectrode array. Differential recording between contacts at 150 micron centers strongly attenuates volume conduction from distant sources, thus assuring that activity is locally generated^12,13^. Spectral content of the resulting LFPg was estimated using the Fast Fourier Transform at each cortical depth. Normalizing within each frequency across channels allowed us to measure the relative contribution of each cortical layer to different frequency bands. A striking finding was that delta and theta oscillations (<10Hz) were focally generated in the superficial layers of all 19 subjects and cortical regions during both wakefulness and sleep (Fig. 1 c,d,e) (**Supplementary Fig. 2**) (Wilcoxon Sign Rank test comparing delta/theta power (1-9 Hz) in channels 1-5 vs. 6-23 across subjects, p<.0001). Although delta/theta power in superficial layers (1-9 Hz, channels 1-5) was on average 30% higher during sleep than wakefulness, it was generated in the same cortical layers in both states (Fig. 1 d).

**Fig. 1.**
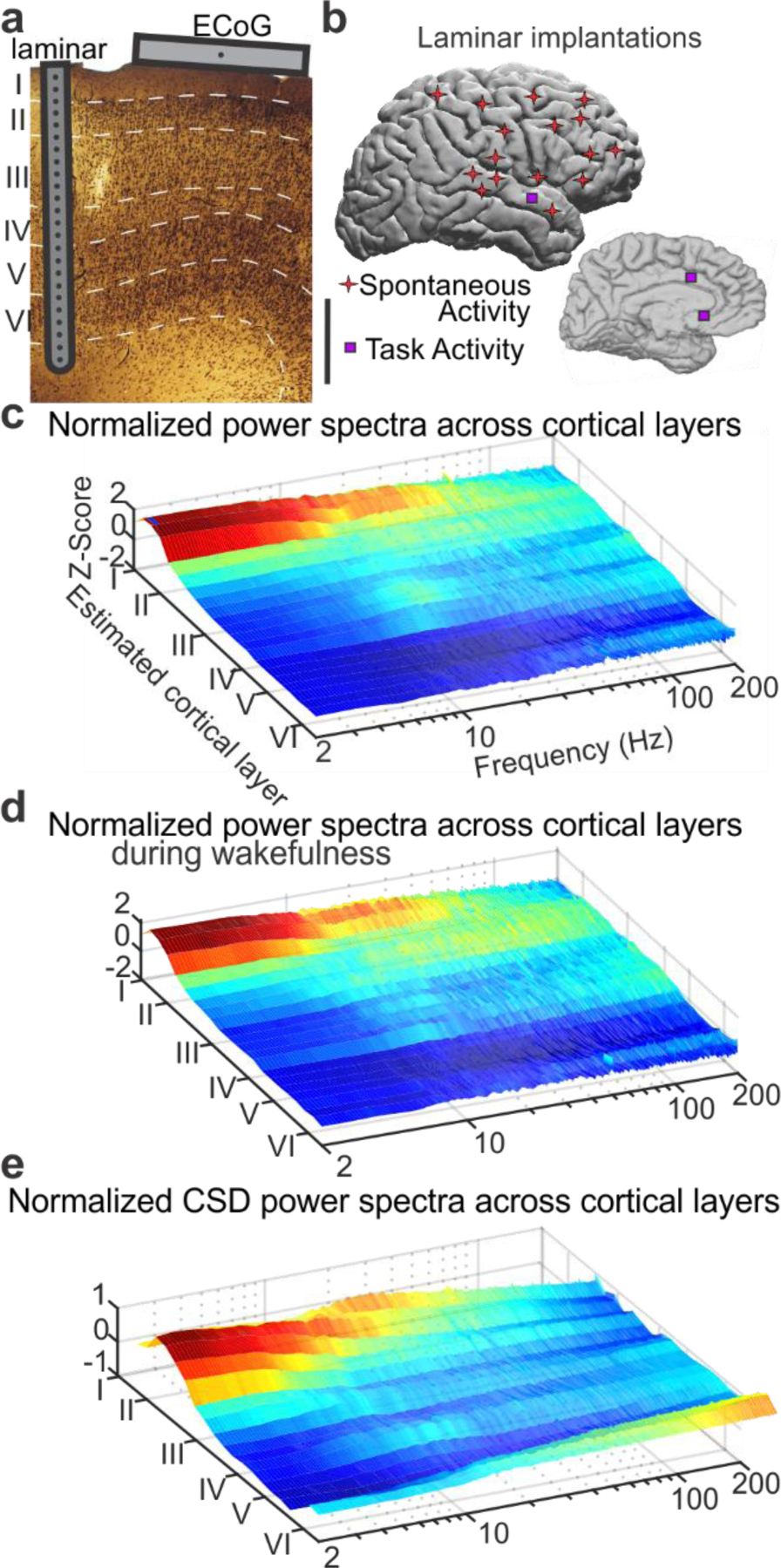
Slow rhythms are generated in superficial cortical layers. (**a**) A schematic of an implanted laminar array and surface ECoG contact in a single patient, overlying a histological section taken from an implantation site. The laminar array comprises 24 contacts on 150µ centers. Each bipolar Local Field Potential gradient (LFPg) recording measures activity from one layer of a single cortical domain. In contrast, the ECoG grid measures activity averaged across all layers from multiple cortical domains (Scale bar: 1mm). The microelectrode reference scheme is illustrated in **Supplementary Fig. 1**. (**b**) Approximate location of laminar implantation in 19 patients. Implants could be in either hemisphere. (**c**) Overall distribution of spectral power across cortical layers. Spontaneous LFPg power was z-normalized across layers to correct for 1/f scaling, and averaged across 16 subjects for sleep and wakefulness, using 1-Hz Gaussian frequency smoothing. Individual subject data is shown in **Supplementary Fig. 2**. (**d**) Normalized power spectral density during wake recordings only. Note that delta/theta band activity is still focally generated in superficial cortical layers. (**e**) Normalized power spectral density using current source density (CSD) instead of the local field potential gradient (LFPg). Delta/theta oscillations are still localized to superficial layers.

Although the concentration of power in superficial layers was most prominent in lower frequencies, it also was present above 10 Hz: 10-100 Hz power in contacts 1-5 (approximate layers I/II) was significantly greater than in contacts 6-23 (Wilcoxon Sign Rank, p<.00044). The only consistent exception to this general principal was the concentration of 10-20Hz power in middle cortical layers (contacts 10-13) noted in several sleep recordings (Fig. 1 c,d,e **Supplementary Fig. 2**). This may reflect spindle generators in thalamorecipient layer IV^14,15^.

### Coherence

We determined if these superficial delta/theta oscillations were synchronized across the cortical surface by measuring their coherence (phase and amplitude consistency) between pairs of ECoG macroelectrode contacts (Fig. 2a). Extending previous findings^16^, we found that delta and theta band oscillations were highly and significantly coherent across the cortical surface, whereas faster oscillations had progressively less lateral spread during both sleep and wakefulness (Fig. 2 A, **Supplementary Fig. 4a**). Measuring coherence across cortical layers demonstrated a similar profile of maximal synchrony at short distances and slow frequencies (Fig. 2b, **Supplementary Fig. 4b**) suggesting that slower rhythms mediate spatial integration both across areas and across layers. Although interlayer coherence varied from subject to subject, presumably reflecting differences in the local cortical microcircuits of various areas^17^, it was consistently significant and maximal in very superficial channels (approximate cortical layers I-II) in the delta/theta band (Fig. 2c, **Supplementary Fig. 3c, Supplementary Fig. 4c**).

**Fig. 2.**
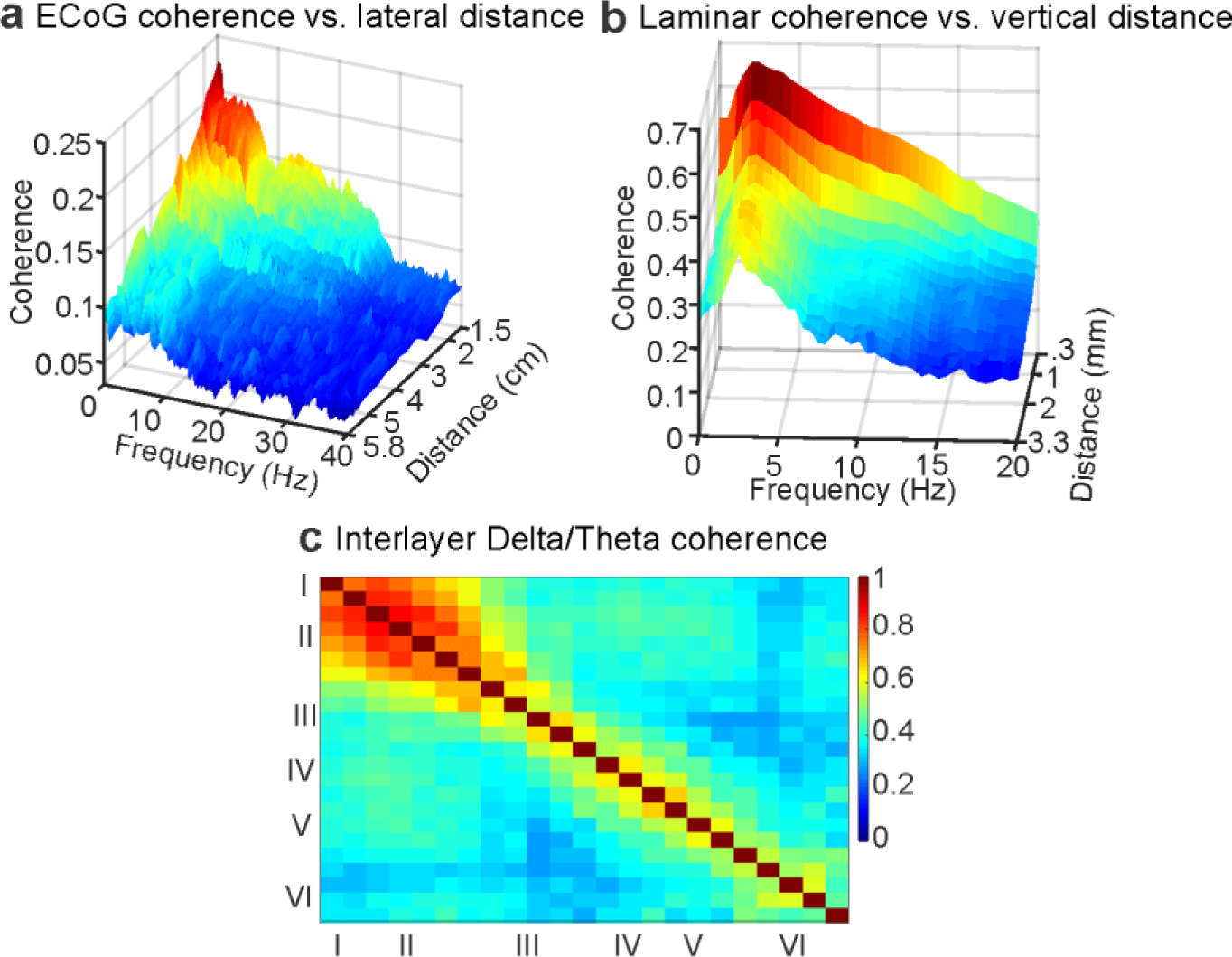
Superficial slow rhythms are coherent across cortical layers and areas. (**a**) Coherence as a function of frequency and distance between cortical regions, measured with bipolar ECOG derivations, and averaged across 4 subjects. (**b**) Coherence as a function of frequency and distance between cortical layers, measured with bipolar laminar derivations. (**c**) Coherence between specific layers in delta/theta band. Individual subject data is shown in **Supplementary Fig. 3 B**. B-C are grand averages across 16 subjects. Significance by distance, frequency and layer for panels a, b and c is shown in **Supplementary Fig. 4**.

We hypothesized that the high amplitude delta/theta in superficial cortical layers generated the laterally coherent rhythms measured with ECoG. This was tested with a multi-scale approach by measuring the coherence between simultaneously recorded laminar and ECoG contacts, allowing us to determine which cortical depth and frequency was most coherent with contacts on the cortical surface (Fig. 3a, b). Confirming our hypothesis, high delta/theta coherence was found between superficial laminar contacts and distant ECoG contacts in both sleep and wake states, maximal in contacts close to the laminar electrode (Fig. 3b, **Supplementary Fig. 5, Supplementary Fig. 6a**). In one subject, we measured the coherence between a depth electrode in the hippocampus and the laminar microelectrode array. Superficial cortical activity was significantly coherent with hippocampal activity in the same delta and theta bands (Fig. 3d, e, **Supplementary Fig. 6b**), confirming that delta/theta rhythms synchronize hippocampal activity with other brain areas^18^.

**Fig. 3.**
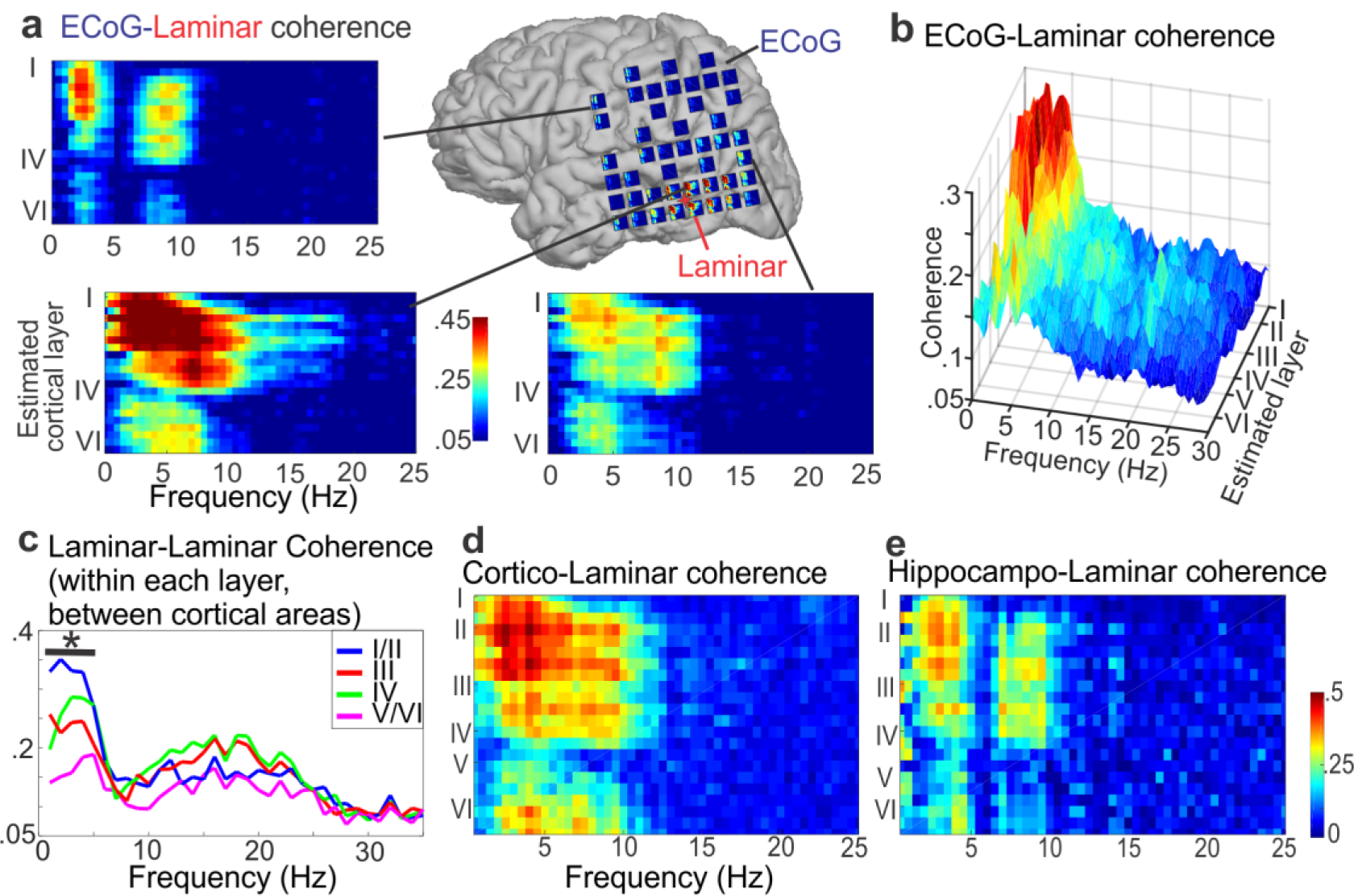
Slow rhythms in superficial cortical layers are coherent with other cortical areas. (**a**) Coherence (frequency vs. cortical depth) of individual ECoG contacts with the laminar probe in a single awake participant. There are high levels of delta/theta coherence in contacts as far as primary motor cortex. ECoG contact spacing is 1cm. Data from 3 additional subjects is shown in **Supplementary Fig. 5**. (**b**) Average coherence (n=4) between a laminar probe and all ECoG (surface) contacts as a function of depth and frequency. Significant ECoG-Laminar coherence was found most consistently within low-frequencies and superficial layers. (**c**) Average coherence (n=3) within layers between two simultaneously recorded laminar arrays. The asterisk indicates that coherence between very superficial contacts (approximate layers I/II) was significant within all 3 subjects from 1-5 Hz. (**d**) Coherence between the neocortical laminar array and an example cortical SEEG bipolar macroelectrode. (**e**) Coherence between the laminar array and a bipolar macroelectrode recording within the head of the hippocampus. Single subject. Statistical comparisons for panels b, d and e are shown in **Supplementary Fig. 6**. In all cases, coherence is significant at low frequencies in upper layers.

Although we found high coherence between superficial slow rhythms measured by laminar electrodes and widespread slow rhythms measured by ECoG, this doesn’t directly demonstrate that delta/theta activity is synchronous within the superficial layers. Simultaneous recordings from two laminar probes spaced one centimeter apart in each of three patients allowed us to measure the coherence within and between layers of distinct cortical regions. Consistent with the laminar-ECoG results, coherence was maximal and significant between the superficial layers of both areas in the delta and low theta bands (Fig. 3c).

### Phase-Amplitude Coupling

To gain further insight into the possible role of superficial slow rhythms in cortical processing, we used Tort’s Modulation Index^19^ to investigate whether the phase of slow rhythms in superficial layers modulates the power of faster frequencies in other layers (Fig. 4a-b, **Supplementary Fig. 7**). Delta phase robustly modulated theta power, with an increase in theta-band power during the falling phase of the ongoing delta rhythm (0 to π radians). Both delta and theta oscillations in superficial layers modulated the power of alpha, beta and gamma oscillations during both sleep and wake states, and the power of these faster oscillations were maximal during the up phase of the spontaneous delta/theta oscillation consistent with previous reports^20^.

**Fig. 4.**
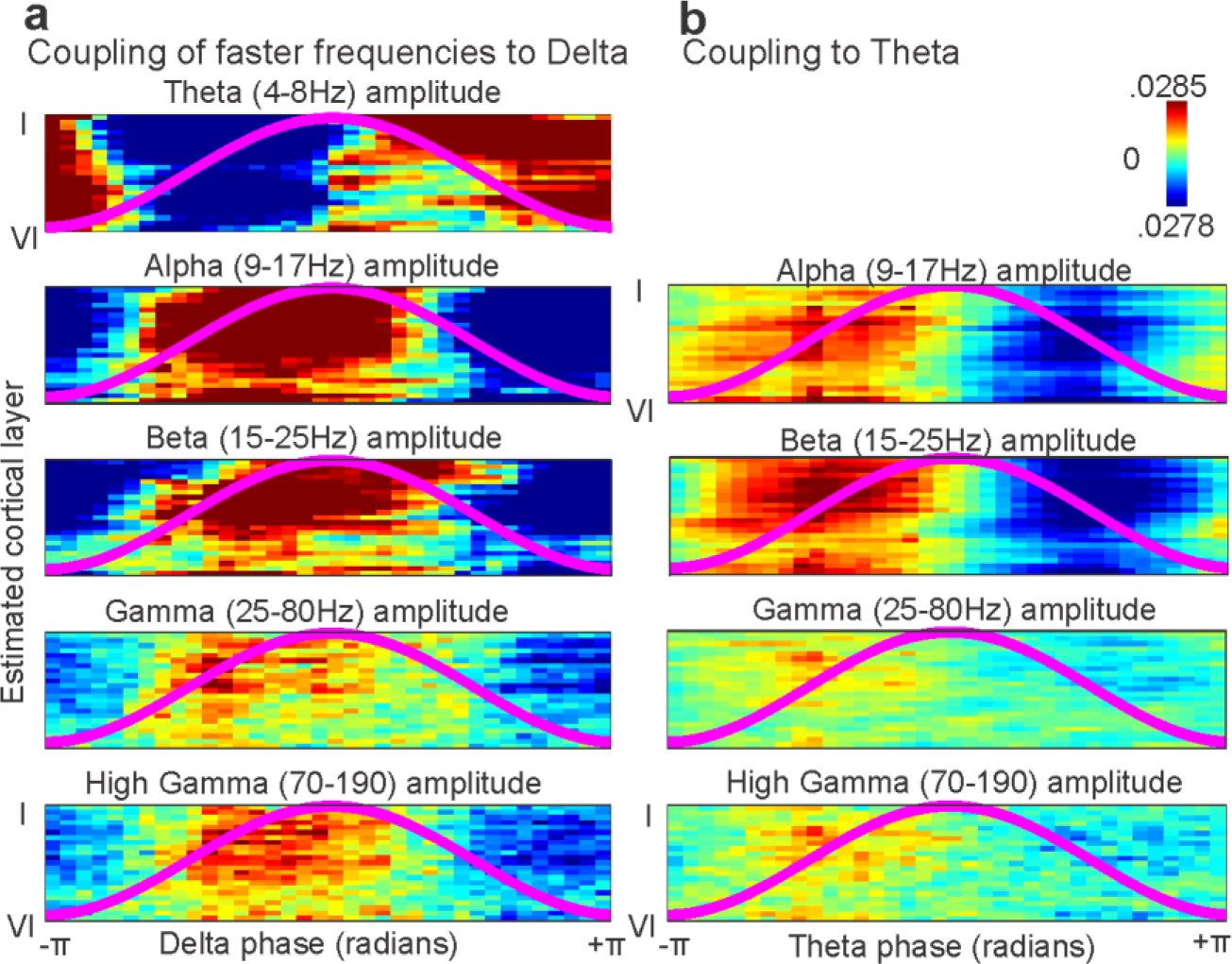
Superficial slow oscillations are coupled to faster frequencies across all layers. (**a**) Phase of ongoing delta in superficial layers is tightly coupled to the amplitude of faster frequencies in all layers. Amplitude was normalized within each contact and frequency band, and then averaged across all subjects and states. Each panel represents phase on the X-axis, cortical layer on the Y-axis and average amplitude of a given frequency in color. The overlying line shows delta phase. (**b**) Same as A, but for coupling of superficial theta phase to amplitude of higher frequencies in all layers.

### Modulation by Cognitive Task

To determine if superficial slow rhythms are modulated by a cognitive task, we used a standard auditory oddball paradigm in which a series of frequent tones, infrequent target tones, and infrequent novel sounds (the latter two referred to as infrequent tones) were presented to 3 participants. We found an evoked response to infrequent stimuli consistently in superficial laminae – the latency, triphasic waveform and frequency (~2 Hz) of the response (Fig. 5a) as well as single trial data (Fig. 5c), leading us to test if the response reflected a phase reset of ongoing slow activity using inter-trial phase clustering (ITPC) in the delta/theta band triggered by stimulus presentation. In two subjects (with laminar probes in the cingulate gyrus) significant ITPC differences were found between frequent and target tones (but not between frequent and novel stimuli), and in a third (laminar probe in the superior temporal gyrus) a significant difference was found between frequent tones and novel auditory stimuli (but not between frequents and infrequent target tones) (p < .05, Bonferroni corrected at each channel by time point) (Fig. 5b). This may reflect areal differences in cognitive function. Delta amplitude was not significantly different between conditions in the cingulate leads. In the temporal lead, a significant increase in delta power did occur mostly in deep cortex, but the increase in delta ITPC as well as the time-domain response were predominantly supragranular. This is consistent with the evoked response being due to a phase-reset rather than an additive response^21,22^. Complete single subject results and significances are plotted in **Supplementary Fig. 8**.

**Fig. 5.**
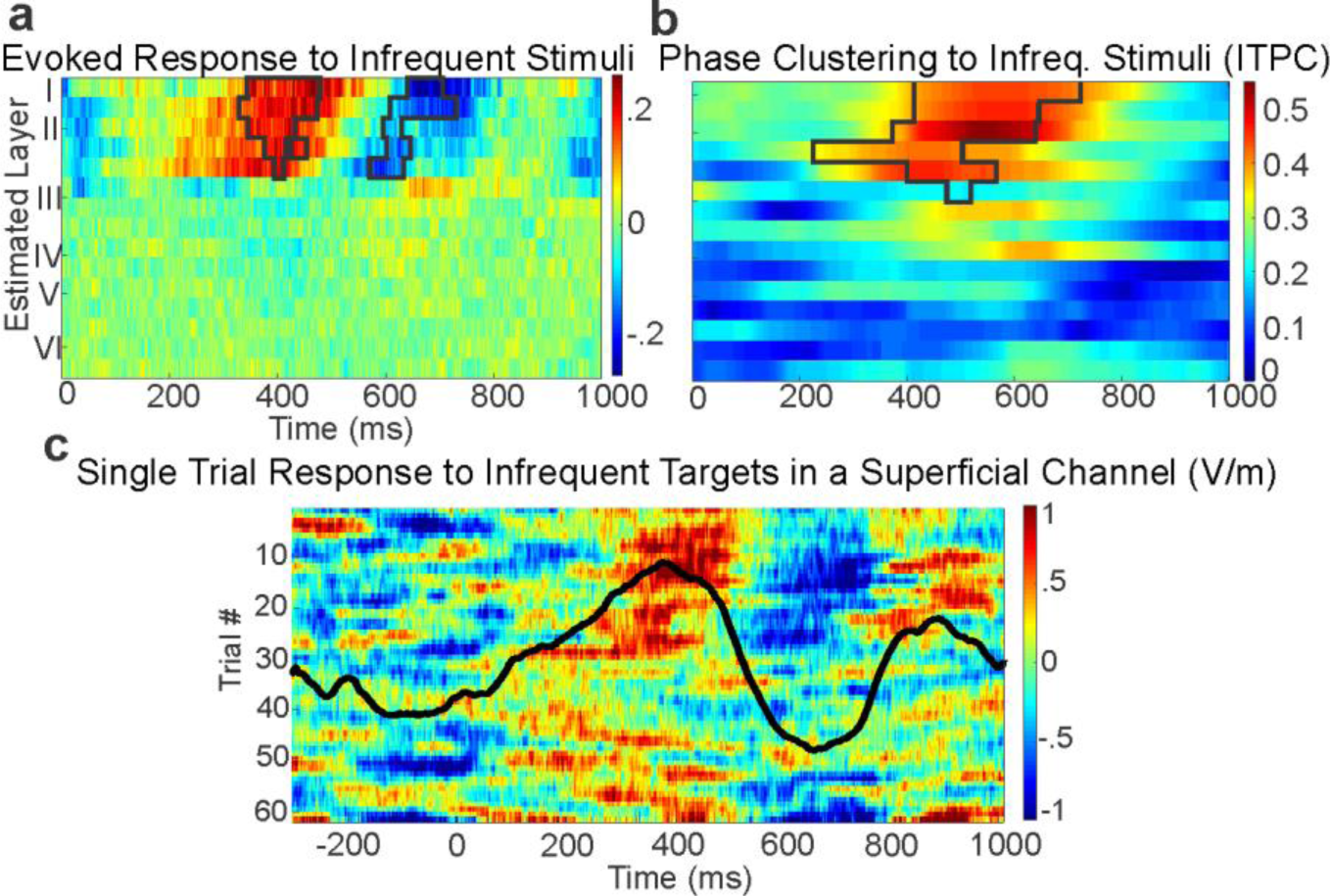
Superficial slow rhythms are phase-reset by infrequent stimuli in an auditory oddball task. **(a**) The average time-domain difference of the LFPg response to infrequent and frequent stimuli (infrequent-frequent) reveals a significant difference in superficial layers at ~400-800 ms after the stimulus in a representative subject. Black lines outline the cluster of channels and time points in which this effect was significant (Nonparametric permutation cluster test, p<.01) (**b**) Slow activity (<3 Hz) is phase locked by infrequent stimuli in superficial channels. As in Fig. 5a, lines border the significant cluster (Nonparametric permutation cluster test, p < .01). (**c**) Single-trial raw data from a superficial channel indicates that infrequent stimuli reset the phase of an ongoing slow oscillation; the channel’s time domain average is overlaid.

## Discussion

We used multiscale recordings to define the distribution and coherence of field potential generation across different cortical layers and areas in humans. A consistent finding across all 19 subjects was that slow (delta/theta) rhythms were generated in and coherent throughout the superficial layers of the cortex, and were coupled to high frequency activity in all layers. In three subject, these rhythms were found to be reset by infrequent stimuli. These findings have practical implications for the neural basis of extracranial EEG as well as the physiology of the cortex and the integration of neural activity.

EEG is a mainstay of clinical neurology and the most widely used method for monitoring brain activity with millisecond precision^23,24^. Our results shed light on its genesis, suggesting that the human EEG consists mainly of relatively slow and coherent activity generated in the superficial cortical lamina correlated with presumed firing in all layers.

The strength of a given cortical layer’s contribution to the scalp EEG depends on the a) power and b) coherence of its currents throughout the cortical mantle^25^. Our study used microelectrode recordings to characterize the power of EEG activity at different cortical depths and frequencies, and a combination of micro and macroelectrodes to characterize its coherence. Laminar array recordings which densely sampled all cortical layers indicated that over 70% of LFPg power is below 10Hz, and is generated in the upper 20% of the cortex. This finding is in contrast to some prior laminar recording studies which emphasized deep sources^26^. However, those studies used referential recordings in which deep channels were contaminated by volume conduction from more superficial sources^13,27^.

Furthermore, we demonstrated that the slow rhythms measured in a single cortical location with a laminar probe or ECoG contact were coherent throughout superficial layers. This was proven indirectly by measuring the coherence between ECoG recordings and laminar LFPg, then directly by measuring the coherence between two simultaneously recorded laminar probes. In both cases, superficial slow LFPg was significantly coherent throughout the cortex. Thus, the slow waves generated across the cortical surface would be expected to summate in propagating extracranially^25^. In summary, laminar and ECoG recordings provide synergistic evidence for high amplitude, coherent delta/theta activity in superficial layers generating much of the spontaneous scalp EEG.

The concentration of delta/theta activity in upper layers was present across all 19 subjects, in parietal, frontal and temporal lobes, in both hemispheres, and generalized across behavioral state and frequencies. While only associative cortex was sampled here, some results in animals suggest that these rhythms extend to primary sensory areas (^13,20^). They may also be more prominent in associative cortex due to its relaltively slower processing timescale^28,29^ and in higher primates because of their expanded supragranular cortex^30^. Although our focus was on delta, and more of our recordings were from sleep than waking when these frequencies have maximum power, we found a similar concentration of power in upper layers during wakefulness. The presence of waking delta is consistent with previous iEEG reports^31^.

Although the dominance of superficial activity throughout the cortex may lead one to posit that this is a biophysical effect (and not due to neural activity per se), we can find no plausible biophysical factors which could explain our results. While differences in impedance could cause systematic laminar biophysical differences in the LFPg, impedance spectra are uniform across cortical layers^32^. Differences in dendritic diameter could also cause differences in the LFPg between layers, as apical dendritic shafts can act as low-pass filter^33^. However, low-pass filtering would only explain differences in high frequency power between layers, and not the concentration of delta/theta in superficial cortex. Furthermore, similar results were found using current-source-density analysis. These considerations suggest that our finding that low frequency LFPg is largest in superficial layers reflects its local generation, rather than biophysical factors.

Our analysis was not confined to sinusoidal oscillations but included all spontaneous activity in the recorded epochs. It may be that specific oscillatory trains which are clearly distinguished from background activity could be concentrated in layers other than superficial, and indeed some sleep recordings displayed what appeared to be spindle activity maximal in middle layers. However, consistent with our findings, previous work has found predominantly superficial CSD activity underlying spontaneous large single^34^ or repeated^2,22^ waves in the delta/theta band.

Superficial slow activity was preferentially phase locked to unexpected sounds, consistent with these rhythms consolidating stimuli into a global cognitive context. Importantly, delta power did not increase relative to infrequent stimuli, demonstrating that the evoked response was due to the phase-reset of ongoing delta oscillations rather than a separate potential. The laminar distribution of delta/theta activity during the task was the same as that of spontaneous slow rhythms, as had been noted previously^22^. The latency and waveform of the evoked response generated by this delta/theta phase reset suggests that superficial slow rhythms contribute to the P3b, associated with cognitive integration across multiple associative cortical regions^35,36^. These results are consistent with CSD studies which found that later activity influenced by cognitive variables was largely superficial^37^, as well as previous recordings of task-related delta activity in iEEG^38,39^.

The transmembrane currents underlying superficial delta/theta LFPg may arise from either voltage-gated or ligand-gated channels^40^. One plausible voltage-gated candidate are H currents, which are most known for contributing to spindle activity but can also generate lower frequency oscillations^41^. Interestingly, the density of H-currents on the dendrites of layer Vb pyramidal cells has been found to increase dramatically in more superficial layers^42^. Alternatively, the slow rhythms we observed may be generated by ligand-gated receptors such as NMDA and GABA-B receptors, which have the long timescales necessary to generate slow activity and are also predominantly in superficial cortical layers^43^. On the circuit level, thalamocortical matrix afferents terminate primarily in upper layers and could mediate the transcortical coherence we observed via their diffuse, modulatory projections^10,44^. In addition, cortico-cortical association fibers terminate mainly in upper layers and would also be expected to contribute to coherence between areas^7–9,45^.

We also demonstrated that superficial delta/theta LFPg was tightly phase-coupled to high gamma power throughout the cortical depth. Since high gamma reflects unit firing^46^, this suggests that the delta/theta LFPg reflects an intracortical circuit engaging cells in all layers, even though the principal generating currents are superficial. These findings extend previous studies which found theta-high gamma coupling in surface ECOG^47^ and general embedding of higher frequencies by lower in laminar recordings in monkeys^20^. Although no strong return current source was observed in deep layers, it is nonetheless likely that currents in the apical dendrites of deep as well as superficial pyramidal cells contribute to the superficial generating sink. The deep source may be attenuated by spatial dispersion resulting from the involvement of both supragranular and infragranular pyramids in the superficial sink, as well as even more superficial sources^2^.

The coherence of the superficial delta/theta across both the depth and extent of the cortex suggests that it plays a central role in the integration of cortical processing. Consistent with this interpretation, the various mechanisms cited above as potential generators of superficial delta/theta such as matrix thalamocortical and top-down cortico-cortical fibers as well as NMDA and GABA-B synapses have been posited to support this integration. Generally, higher associative cortico-cortical feedback processes are thought to have relatively slow timescales^3–6^, and to be primarily supported by superficial cortex^7–10,45^. Our data are consistent with a circuit model of cortical integration in which endogenous feedback inputs onto the apical dendrites of pyramidal cells modulate feedforward activity and neuronal firing in deeper layers^9,48^, as well as a model in which deep fast activity plays a role in regulating superficial slow rhythms^49^. In this view, the EEG is mainly generated by superficial slow currents which are the summated signal of widespread cortical associations.

## Methods

### Participants

Nineteen participants (5 female, ages 12-55) with pharmacologically resistant epilepsy were implanted with intracranial electrodes to localize seizure foci. All 19 subjects had at least one laminar probe. We analyzed an average of 17.2 minutes of clean, spontaneous activity from 16/19 participants and task activity from 3/19. These implantations were performed with fully informed consent as specified by the Declaration of Helsinki and approved by local institutional review boards. These review boards were the Partners Health Care IRB, Committee on Clinical Investigations of the Beth Israel Deaconess Medical Center, NYU Medical Center IRB, and the Hungarian Medical Scientific Council. All experiments were performed in accordance with the guidelines and regulations of these review boards. All decisions concerning macroelectrode placement (both surface and depth (sEEG) electrodes) were made solely on a clinical basis. Patients were fully informed of potential risks and were told that they had no obligation to participate in the study, and that their choice not to participate would not affect their clinical care in any way.

### Electrodes

To sample from each cortical layer simultaneously, each laminar array had 24 contacts with 40 µm diameters and 150 µm center-to-center spacing. Each laminar probe spanned the cortical depth with a length of 3.5 mm and diameter of 0.35 mm (Fig. 1a, b) ^11^. Laminar electrodes were of two kinds. The more common ‘surface’ laminar arrays (17 of 19 patients) were inserted perpendicular to the cortical surface under visual control. In order to consistently position the surface laminar arrays relative to cortical layers, a silicone sheet was attached perpendicular to the top of the array anchoring it to the cortical surface. Sheet position was maintained by surface tension and the overlying ECoG array and dura. Thus, physical constraints resulted in the first contact being centered ~150µ below the pial surface, and the 24^th^ contact at ~3600µ below the pial surface. The correspondence of channels to layers was extrapolated from previous measurements of laminar width in human cortex ^30^. Channels 1, 4, 9, 14, 16 and 21 were the approximate centers of layers I-VI, respectively.

The other type of laminar electrode, the ‘depth’ laminar (2 of 19 patients), was inserted through the lumen of the clinical depth electrodes so as to extend past the clinical tip by ~5mm, and the clinical electrodes were implanted ~5mm less deep than they would otherwise have been. Thus, their placement with respect to cortical laminae was less certain, based upon co-registered MRI/CT and confirmed with basic physiological measures, notably the presence of high gamma and/or multiunit activity which is confined to gray matter. Physical constraints determined whether the lead approached the cortical ribbon from the white matter or the pia. In 2 patients with surface laminars, it was possible to confirm placement using histology. One such patient (Subject 14) is shown in Figure 1A. In this patient, delta/theta power was concentrated in supragranular cortex, more specifically layers I/II (contacts 1-5), as was seen in all other patients (**Supplementary Fig. 2**).

Macroelectrodes included electrocorticographic (ECoG) grids (2 mm diameter, 1 cm pitch) and depth electrodes, also known as stereo-EEG (SEEG). The signals were originally recorded with a relatively inactive, clinical reference. This referencing scheme was used for calculating coherence between the laminar array and ECoG contacts. Prior to the computation of coherence between ECoG contacts, a bipolar montage was used wherein each channel was referenced to its right-side neighbor (**Supplementary Fig. 1**). Structural MRI or CT with the electrodes in place, aligned with preoperative MRI, was used to identify the position of SEEG and ECoG contacts ^50,51^. Neighboring pairs of contacts in the hippocampal or cortical gray matter were referenced to each other.

### Recordings

Local field potential recordings from the laminar microelectrode arrays were sampled at 2000 Hz with an online low-pass filter of 500 Hz. Each laminar contact was referenced to its neighbor, yielding the potential gradient. Potential gradient (the first spatial derivative) rather than current-source-density (CSD) (the second spatial derivative) was used for several reasons. Previous studies have shown that using this reference scheme largely eliminates volume conduction and provides a very local measure of cortical activity^12^. Due to the high sensitivity of the second spatial derivative to noise, eliminating spurious sources and sinks requires heavily smoothing the signal prior to taking the derivative. This spatial smoothing would attenuate the laminar precision that CSD is intended to reveal. Furthermore, because the second spatial derivative is estimated using a three to five point approximation, a single faulty contact could necessitate the removal of 3 or more signals, a significant amount of the laminar depth. Lastly, modeling and empirical studies have shown that the potential gradient and CSD both yield similar spatial localization^12,13^. To ensure that our findings were independent of analysis technique, we also found the power spectral density at multiple cortical depths using CSD (Fig. 1e) with the five-point approximation^11^. This yielded very similar results to the potential gradient.

### Epoch selection

All data were visually inspected for movement, pulsation and machine artifacts. The data was also screened for epileptic activity such as interictal discharges and pathological delta by a board certified electroencephalographer. Laminar arrays with significant amounts of artifactual or epileptiform activity, and/or insufficient technical quality, were rejected prior to further analysis. All epochs with artefactual or epileptiform activity from accepted arrays were also excluded from analyses. Despite these measures, contamination due to epilepsy was a concern due to placement of the surface arrays in a location that had a high likelihood of being subsequently resected. Because depth laminars were integrated with the clinical probes they could be placed in locations which were suspected of being involved in the seizure but with less certainty than the surface laminars.

Thus, as a further check, we compared the strength of superficial delta/theta oscillations in spontaneous activity recorded from laminar probes whose recordings exhibited interictal spikes (N=6) versus laminar probes which did not record any interictal spikes in the examined epochs (N=10). Restricting analysis to the epochs without interictal spikes in all participants, we found no significant difference in normalized delta/theta power (Wilcoxon Rank Sum, p=.64); in fact, mean delta/theta power differed between groups by <1%, being slightly higher in healthier electrodes (interictal: 1.1235±.3037, non-interictal: 1.1327±.2270, mean±standard deviation), suggesting that these oscillations are not pathological in nature.

Another source of concern related to epilepsy is the possible effects of Anti-Epileptic Drugs (AED) on the local field potentials reported here, inasmuch as such effects have been reported in the scalp EEG^52^. While some patients in this study may have been taking AEDs, most recordings were performed after the patient’s medications had been tapered to encourage spontaneous seizure occurrences during the monitoring period. More importantly, as described, the results were highly consistent across all participants regardless of medication history, etiology, electrode location, or degree of epileptic activity. Expert screening, consistency across participants, and convergent results in healthy animals^13^ strongly suggest that our findings are generalizable to the non-epileptic population.

### Spectral Analysis

All analysis was performed in MATLAB using custom and FieldTrip functions^53^. In each participant, the Fourier Transform was calculated in 10 second epochs on the zero-meaned data after a single Hanning taper was applied. The power spectrum was then Z-normalized across channels / within each frequency band in order to determine the relative power of different oscillations in different layers. The normalized frequency spectra of faulty channels (an average of 3 per probe) were linearly interpolated from the normalized frequency spectra of good channels above and below on the laminar probe. For instance, if channel 2 was defective, its power spectrum would be replaced by the average of channels 1 and 3’s power spectra, However, note that subjects 2, 3, 7, 8, 9 and 16 (depicted in **Supplementary Fig. 2**) had no interpolated channels yet still had delta/theta activity concentrated in superficial cortex. Linear interpolation of bad channels in the time domain prior to the FFT yielded artificially low high frequency power due to phase cancellation.

### Coherence

The coherence between zero-meaned time series x and y was defined as 
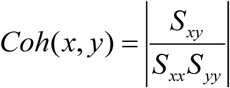
, where *S_xy_* is the cross-spectral density between x and y and *S*_*xx*_ is the autospectral density of x. Because small numbers of epochs can lead to spurious coherence, recorded epochs were subsampled into two second epochs prior to the calculation of auto / cross spectra^23,54^.

To find the statistical significance of coherence values (Fig. 2, Fig. 3), we used a non-parametric trial shuffling procedure^23^. Within each subject, we shuffled the temporal order of two second epochs for each channel 100 times and for each shuffle recomputed all coherencies. We used this distribution of coherencies, generated under the null hypothesis of no temporal relationship between channels, to z-score the real coherencies of each channel pair, giving us single subject z-scores for each pair of contacts and frequency. Unless otherwise specified, coherence was deemed significant if its p-value was < .05, Bonferroni Corrected. More specifically, we set the critical value at .05 / the number of channel by channel combinations within each subject. For instance, if a given subject had 23 functioning laminar contacts, significance would be set at .05 / ((23*22)/2). This procedure was applied within subjects, and then the consistency of the effect across subjects was plotted as the proportion of subjects which had significant coherence for each channel pair and frequency (**Supplementary Fig. 4, Supplementary Fig. 6**).

To determine how coherence varies across the cortical surface or between ECoG contacts, pairs of contacts were sorted by the Euclidean distance between them on the pial surface. Then, the average coherence was found over all pairs of contacts at a given distance. Due to varying intercontact distances present in each subject’s electrode configurations, the average coherence vs. distance matrix for each participant (not the raw time-domain data) was linearly interpolated at every .2 cm before averaging across participants. To determine how coherence varied perpendicular to the cortical surface, the same procedure was applied to the laminar microelectrode array without interpolation.

To assess the significance of average coherence at various intercontact distances and frequencies (Fig. 2), we iteratively computed the average coherence vs. distance map as described above (within subjects) after shuffling trials, and then used these maps (created under a null hypothesis of no temporal relationship between channels) to z-score the real coherence vs. distance map rather than the coherencies per se (**Supplementary Fig. 5**).

Simultaneous ECoG and Laminar recordings were available in four participants, two made from awake participants and two from sleeping participants. The coherence was measured between each pair of ECoG and laminar contacts. The average coherence between each laminar contact and the 20 closest grid contacts was used for plotting and statistical testing. All recordings showed high coherence between grid and superficial laminar contacts in the delta/theta band.

In one participant, coherence was calculated between a bipolar referenced SEEG lead within the hippocampal head and a simultaneously recorded laminar array. The coherence between each laminar signal and the bipolar referenced hippocampal lead was compared to the average coherence between the laminar array and an SEEG lead with two contacts in temporal neocortex (Fig. 3 C, D). Although cortico-cortical coherence is higher in magnitude, both are significantly coherent within the same bands.

In three participants, two laminar arrays spaced one centimeter apart were recorded from simultaneously. One of these participants was awake during the recording, two were asleep. The coherence between each pair of laminar contacts was calculated, and then averaged within putative cortical layers. All participants displayed significant coherence in superficial contacts and low frequencies.

### Phase – Amplitude – Coupling

To determine the effects of superficial slow rhythms on cortical activity, we used Tort’s Modulation Index^19^ with a non-parametric trial shuffling procedure to assess significance. First, the data was split into two second epochs. Then, each trial was filtered within the frequency bands of interest using a fourth-order IIR Butterworth Filter. The first and last 100 ms of data were removed to eliminate edge artifacts. Then, the analytic signal *z*(*t*) was found by applying the Hilbert Transform to the Local Field Potential gradient (LFPg) of each channel. In each recording, the contact with the highest power in the modulating / lower frequency band was used as the ‘phase index’ for determining modulation of power in other channels. All such contacts were within the 5 closest to the laminar entry (i.e. superficial channels). Only epochs with high Hilbert amplitude values for the modulating frequency (top 50%) were analyzed further. The phase series *ϕ*(*t*) of the phase index channel was found by taking the angle of the analytic signal, and the amplitude *A*(*t*) of every channel was found by taking the real component of the analytic signal. *ϕ*(*t*) was then reordered from −*π* to +*π*, and *A*(*t*) for every other channel and frequency was reordered using the same permutation vector. Amplitude was then averaged within 36 bins of phase (i.e. 10 degrees) and normalized by the sum over bins, yielding *ϕ*. The modulation index (MI) was then calculated as 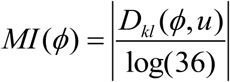 for each channel and frequency pair, where *D_kl_* is the Kullback-Leibler divergence, *u* is the uniform distribution (i.e. no relationship between amplitude and phase) and log(36) is the natural logarithm of the number of phase bins^19^. *D_kl_* was computed as log(36) – H(P), where H(P) was the distribution’s Shannon’s Entropy.

To determine the statistical significance of these MI values, we generated a reference distribution under the null hypothesis of a random relationship between amplitude and phase by iteratively shuffling the phase series (by splitting the phase series into two epochs and swapping their order) and recalculating the MI for each shuffled dataset. The mean and variance of these null hypothesis derived MIs at each channel and frequency were used to determine the z-score of the actual MIs at each channel and modulating/modulated frequency pair, with significance set at p < .05, Bonferroni Corrected (the critical value was set at .05 / 14, 14 being number of modulating/modulated frequency pairs, within each subject). The percentage of subjects with a significant MI between each pair of channels and modulating/modulated frequencies was then plotted within states (**Supplementary Fig. 7**). This analysis indicated a significant modulation of high frequency amplitude by delta and theta-phase throughout the cortical depth within subjects as well as states (p < .05, Bonferroni Corrected).

### Auditory Oddball Task

A standard auditory oddball paradigm allowed us to assess the cognitive correlates of superficial slow activity. Stimuli consisted of frequent (80%) tones, infrequent target (10%) tones and infrequent novel (10%) sounds with a stimulus onset asynchrony of 1600 ms. High (600 Hz) or low (140 Hz) tones as targets were counterbalanced across blocks. The participant was asked to silently count and report the number of target tones in each block. To determine significance, a nonparametric permutation cluster test was applied to the differences in the time-domain, delta/theta analytic amplitude and inter-trial phase clustering (ITPC) between frequent tones and novel sounds, as well as between frequent tones and infrequent tones^23^. The same procedure was applied to find significant differences in ITPC, delta amplitude and the LFPg. First, a reference distribution for each channel and time point under the null hypothesis was estimated by shuffling the trial labels between each condition 500 times, and then calculating the average difference between (shuffled) target and filler trials at each channel x time point. This yielded the mean and standard deviation of inter-condition differences under the null hypothesis of no difference between conditions. The actual difference between conditions was z-scored using this non-parametric reference distribution. To correct for multiple comparisons, we z-scored each matrix of differences between shuffled conditions and recorded the size of each contiguous cluster of channel x time points which had p<.05. This yields a distribution of cluster sizes of significantly different points under the null hypothesis, and the 99^th^ percentile of this set of cluster sizes was used as a threshold for cluster significance. Finally, this cluster size threshold was then applied to the original, real Z-scored difference matrix of channel x time points between conditions to determine significance corrected for multiple comparisons^23^.

For ITPC, the entire dataset was filtered from 1-3 Hz, as the time-domain response had a period of ~500 ms. To calculate the inter-trial phase clustering value (ITPC) and determine if the P3 stemmed from a phase reset, we then applied the Hilbert Transform to our dataset and took its angle to find the instantaneous delta phase at each point in time. The phase at each channel and time point was then represented as a complex vector in the unit circle using Euler’s Identity formula *e^iθ^* where *θ* is delta phase. The ITPC was then calculated as the length of the average phase vector across trials, yielding an ITPC value bound by 0 (no consistency) and 1 (perfect consistency) ^23^. To determine whether or not there was a difference in delta amplitude between conditions, we found the Hilbert amplitude of the filtered data. Significant differences in ITPC (as assessed with the above nonparametric statistics) were found between frequent tones and novel sounds in one subject, and between frequent and infrequent tones in two others^23^.

For visualization of the single-trial data (Fig. 5c) a 10-trial Gaussian smoothing window was applied. A 40-ms wide Gaussian (σ = 4 ms) smoothing of the overlaid time-domain response was also plotted.

### Data Availability

Data and code will be made available upon reasonable request to the degree it is possible given participant consent constraints and HIPAA requirements.

## Acknowledgements

The authors thank Terrence Sejnowski, Erica Johnson, Emília Tóth, Lopes Da Silva, Arnold Mandell, Bjorn Merker, J.F. Bartscher, Qianqian Deng, ChunMao Wang, Adam Niese, and Ksenija Marinković for commentary, feedback and/or technical support. This work was supported by the U.S. Office of Naval Research Grant N00014-13-1-0672, National Institutes of Health Grants R01-MH-099645, R01-EB-009282, R01-NS-062092, K24-NS-088568, the MGH Executive Council on Research, Hungarian National Brain Research Program grant KTIA_13_NAP-A-IV/1-4,6, EU FP7 600925 NeuroSeeker, and Hungarian Government grants KTIA-NAP 13-1-2013-0001, OTKA PD101754.

## Author Contributions

S.C., E.H. and I.U. conceived of the study. I.U., J.M., L.E., S.C., E.H., D.F., O.D. and W.K.D. collected the data. M.H. performed the analysis. M.H., S.C. and E.H. wrote the paper. All authors discussed and edited the manuscript.

## References

1. Buzsáki, G. & Draguhn, A. Neuronal Oscillations in Cortical Networks. Sci. 304, 1926–1926 (2004).

2. Csercsa, R. et al. Laminar analysis of slow wave activity in humans. Brain 133, 2814–2814 (2010).

3. Destexhe, A., Contreras, D. & Steriade, M. Spatiotemporal Analysis of Local Field Potentials and Unit Discharges in Cat Cerebral Cortex during Natural Wake and Sleep States. J. Neurosci. 19, 4595–4595 (1999).

4. Von Stein, A. & Sarnthein, J. Different frequencies for different scales of cortical integration: From local gamma to long range alpha/theta synchronization. Int. J. Psychophysiol. 38, 301–301 (2000).

5. Buffalo, E. a, Fries, P., Landman, R., Buschman, T. J. & Desimone, R. Laminar differences in gamma and alpha coherence in the ventral stream. Proc. Natl. Acad. Sci. U. S. A. 108, 11262–11262 (2011).

6. van Kerkoerle, T. et al. Alpha and gamma oscillations characterize feedback and feedforward processing in monkey visual cortex. Proc. Natl. Acad. Sci. 111, 14332–14332 (2014).

7. Pucak, M., Levitt, J., Lund, J. & Lewis, D. Patterns of intrinsic and associational circuitry in monkey prefrontal cortex. J. Comp. Neurol. 376, 614–614 (1996).

8. Douglas, R. J. & Martin, K. A. C. Neuronal Circuits of the Neocortex. Annu. Rev. Neurosci. 27, 419–419 (2004).

9. Larkum, M. E., Senn, W. & Lüscher, H.-R. Top-down Dendritic Input Increases the Gain of Layer 5 Pyramidal Neurons. Cereb. Cortex 14, 1059–1059 (2004).

10. Rubio-Garrido, P., Pérez-De-Manzo, F., Porrero, C., Galazo, M. J. & Clascá, F. Thalamic input to distal apical dendrites in neocortical layer 1 is massive and highly convergent. Cereb. Cortex 19, 2380–2380 (2009).

11. Ulbert, I., Halgren, E., Heit, G. & Karmos, G. Multiple microelectrode-recording system for human intracortical applications. J. Neurosci. Methods 106, 69–69 (2001).

12. Kajikawa, Y. & Schoeder, E. How local is the local field potential? Neuron 72, 847–847 (2012).

13. Haegens, S. et al. Laminar Profile and Physiology of the α Rhythm in Primary Visual, Auditory, and Somatosensory Regions of Neocortex. J. Neurosci. 35, 14341–14341 (2015).

14. Bonjean, M. et al. Interactions between Core and Matrix Thalamocortical Projections in Human Sleep Spindle Synchronization. J. Neurosci. 32, 5250–5250 (2012).

15. Piantoni, G., Halgren, E. & Cash, S. S. The contribution of thalamocortical core and matrix pathways to sleep spindles. Neural Plasticity 2016, (2016).

16. Arnulfo, G., Hirvonen, J., Nobili, L., Palva, S. & Palva, J. M. Phase and amplitude correlations in resting-state activity in human stereotactical EEG recordings. Neuroimage 112, 114–114 (2015).

17. Ninomiya, T., Dougherty, K., Godlove, D. C., Schall, J. D. & Maier, A. Microcircuitry of agranular frontal cortex: contrasting laminar connectivity between occipital and frontal areas. J. Neurophysiol. 113, 3242–3242 (2015).

18. Fell, J. & Axmacher, N. The role of phase synchronization in memory processes. Nat Rev Neurosci 12, 105–105 (2011).

19. Tort, A. B. L., Komorowski, R., Eichenbaum, H. & Kopell, N. Measuring Phase-Amplitude Coupling Between Neuronal Oscillations of Different Frequencies. J. Neurophysiol. 104, 1195–1195 (2010).

20. Lakatos, P. et al. An oscillatory hierarchy controlling neuronal excitability and stimulus processing in the auditory cortex. J. Neurophysiol. 94, 1904–1904 (2005).

21. Fell, J. et al. Neural Bases of Cognitive ERPs: More than Phase Reset. J. Cogn. Neurosci. 16, 1595–1595 (2004).

22. Halgren, E. et al. Laminar profile of spontaneous and evoked theta: Rhythmic modulation of cortical processing during word integration. Neuropsychologia 76, 108–108 (2015).

23. Cohen, M. X. Analyzing Neural Time Series Data: Theory and Practice. (The MIT Press, 2014).

24. Cohen, M. X. Where Does EEG Come From and What Does It Mean? Trends in Neurosciences 40, 208–208 (2017).

25. Musall, S., von Pfostl, V., Rauch, A., Logothetis, N. K. & Whittingstall, K. Effects of Neural Synchrony on Surface EEG. Cereb. Cortex 24, 1045–1045 (2014).

26. Petsche, H., Pockberger, H. & Rappelsberger, P. On the search for the sources of the electroencephalogram. Neuroscience 11, 1–1 (1984).

27. Kajikawa, Y. & Schroeder, C. E. Generation of field potentials and modulation of their dynamics through volume integration of cortical activity. J. Neurophysiol. 113, 339–339 (2015).

28. Honey, C. J. et al. Slow Cortical Dynamics and the Accumulation of Information over Long Timescales. Neuron 76, 423–423 (2012).

29. Murray, J. D. et al. A hierarchy of intrinsic timescales across primate cortex. Nat. Neurosci. 17, 1661–1661 (2014).

30. Hutsler, J. J., Lee, D.-G. & Porter, K. K. Comparative analysis of cortical layering and supragranular layer enlargement in rodent carnivore and primate species. Brain Res. 1052, 71–71 (2005).

31. Sachdev, R. N. S. et al. Delta rhythm in wakefulness: evidence from intracranial recordings in human beings. J. Neurophysiol. 114, 1248–1248 (2015).

32. Logothetis, N. K., Kayser, C. & Oeltermann, A. In vivo measurement of cortical impedance spectrum in monkeys: implications for signal propagation. Neuron 55, 809–809 (2007).

33. Lindén, H., Pettersen, K. H. & Einevoll, G. T. Intrinsic dendritic filtering gives low-pass power spectra of local field potentials. J. Comput. Neurosci. 29, 423–423 (2010).

34. Cash, S. S. et al. The Human K-Complex Represents an Isolated Cortical Down-State. Science (80-.). 324, 1084–1084 (2009).

35. Halgren, E., Marinkovic, K. & Chauvel, P. Generators of the late cognitive potentials in auditory and visual oddball tasks. Electroencephalogr. Clin. Neurophysiol. 106, 156–156 (1998).

36. Soltani, M. & Knight, R. T. Neural origins of the P300. Critical reviews in neurobiology 14, 199–199 (2000).

37. Wang, C., Ulbert, I., Schomer, D. L., Marinkovic, K. & Halgren, E. Responses of Human Anterior Cingulate Cortex Microdomains to Error Detection, Conflict Monitoring, Stimulus – Response Mapping, Familiarity, and Orienting. J. Neurosci. 25, 604–604 (2005).

38. Nacher, V., Ledberg, a., Deco, G. & Romo, R. Coherent delta-band oscillations between cortical areas correlate with decision making. Proc. Natl. Acad. Sci. 110, 15085–15085 (2013).

39. Szczepanski, S. M. et al. Dynamic Changes in Phase-Amplitude Coupling Facilitate Spatial Attention Control in Fronto-Parietal Cortex. PLoS Biol. 12, e1001936 (2014).

40. Murakami, S., Hirose, A. & Okada, Y. C. Contribution of ionic currents to magnetoencephalography (MEG) and electroencephalography (EEG) signals generated by guinea-pig CA3 slices. J. Physiol. 553, 975–975 (2003).

41. McCormick, D. a & Bal, T. Sleep and arousal: thalamocortical mechanisms. Annu. Rev. Neurosci. 20, 185–185 (1997).

42. Kole, M. H. P., Hallermann, S. & Stuart, G. J. Single I_h_ Channels in Pyramidal Neuron Dendrites: Properties, Distribution, and Impact on Action Potential Output. J. Neurosci. 26, 1677 LP-1687 (2006).

43. Eickhoff, S. B., Rottschy, C. & Zilles, K. Laminar distribution and co-distribution of neurotransmitter receptors in early human visual cortex. Brain Struct. Funct. 212, 255–255 (2007).

44. Jones, E. G. Thalamic circuitry and thalamocortical synchrony. Philos. Trans. R. Soc. B Biol. Sci. 357, 1659–1659 (2002).

45. Burke, S. N. et al. Differential encoding of behavior and spatial context in deep and superficial layers of the neocortex. Neuron 45, 667–667 (2005).

46. Ray, S., Crone, N. E., Niebur, E., Franaszczuk, P. J. & Hsiao, S. S. Neural Correlates of High-Gamma Oscillations (60-200 Hz) in Macaque Local Field Potentials and Their Potential Implications in Electrocorticography. J. Neurosci. 28, 11526–11526 (2008).

47. Canolty, R. T. et al. High Gamma Power Is Phase-Locked to Theta Oscillations in Human Neocortex. Science (80-.). 313, 1626–1626 (2006).

48. Larkum, M. A cellular mechanism for cortical associations: an organizing principle for the cerebral cortex. Trends Neurosci. 36, 141–141 (2013).

49. Jiang, H., Bahramisharif, A., van Gerven, M. A. J. & Jensen, O. Measuring directionality between neuronal oscillations of different frequencies. Neuroimage 118, 359–359 (2015).

50. Yang, A. I. et al. Localization of dense intracranial electrode arrays using magnetic resonance imaging. Neuroimage 63, 157–157 (2012).

51. Dykstra, A. R. et al. Individualized localization and cortical surface-based registration of intracranial electrodes. Neuroimage 59, 3563–3563 (2012).

52. Blume, W. T. Drug effects on EEG. J. Clin. Neurophysiol. 23, 306–306 (2006).

53. Oostenveld, R., Fries, P., Maris, E. & Schoffelen, J.-M. FieldTrip: Open Source Software for Advanced Analysis of MEG, EEG, and Invasive Electrophysiological Data. Comput. Intell. Neurosci. 2011, 1–1 (2011).

54. Rosenberg, J. R., Amjad, A. M., Breeze, P., Brillinger, D. R. & Halliday, D. M. The Fourier approach to the identification of functional coupling between neuronal spike trains. Prog. Biophys. Mol. Biol. 53, 1–1 (1989).

